# Benthic jellyfish dominate water mixing in mangrove ecosystems

**DOI:** 10.1101/784173

**Authors:** David M. Durieux, Kevin T. Du Clos, Brad J. Gemmell

## Abstract

Water mixing is a critical mechanism in marine habitats that governs many important processes, including nutrient transport. Physical mechanisms, such as winds or tides, are primarily responsible for mixing effects in shallow coastal systems, but the sheltered habitats adjacent to mangroves experience very low turbulence and vertical mixing. The significance of biogenic mixing in pelagic habitats has been investigated but remains unclear. In this study we show that the upside-down jellyfish *Cassiopea* sp. plays a significant role with respect to biogenic contributions to water column mixing within its shallow natural habitat (< 2 m deep). The mixing contribution was determined by means of high-resolution flow velocimetry methods in both the laboratory and in the natural environment. We demonstrate that *Cassiopea* sp. continuously pumps water from the benthos upward in a vertical jet with flow velocities on the scale of centimeters per second. The volumetric flow rate was calculated to be 212 l h^−1^ for average sized animals (8.6 cm bell diameter), which translates to turnover of the entire water column every 15 minutes for a median population density (29 animals m^−2^). In addition, we found *Cassiopea* sp. are capable of releasing porewater into the water column at an average rate of 2.64 ml h^−1^ per individual. The release of nutrient-rich benthic porewater combined with strong contributions to water column mixing, suggest a role for *Cassiopea* sp. as an ecosystem engineer in mangrove habitats.

**Significance Statement:** Water mixing is a critical process for aquatic life. Coastal mangrove habitats are vital nurseries for commercially and ecologically important species, but these sheltered habitats experience little water mixing. The upside-down jellyfish, *Cassiopea* sp., occurs in circumtropical mangrove habitats at high densities. They are epibenthic and pulse nearly continuously, producing a vertical current that transports hundreds of liters of seawater per hour. This results in turnover of the entire water column every 15 minutes for an average population. Additionally, *Cassiopea* sp. can greatly expedite the transport of nutrient-rich water from sediments into the water column. Thus, *Cassiopea* sp. represents a previously unrecognized ecosystem engineer that can affect primary productivity, nutrient distribution, and alter new habitats as their range is expanding.

## Introduction

Coastal mangrove habitats exhibit exceptionally high productivity and provide many ecosystem services, including control of flooding, sedimentation, and nutrient input in surrounding areas (1). In addition, these habitats act as critical nursery habitat for early life stages of a wide range of commercially and ecologically important species of fish and crustaceans (2, 3) and as adult habitat for many others (3). Mixing of the water column is a critical process which regulates important processes such as productivity (4) and benthic-pelagic coupling (5), prey encounter rate (6), and gas exchange (7). Thus, it is important to identify and understand the major sources of mixing within these coastal environments. Mixing forces are generally physical in origin, such as wind or tides, but in some cases they can originate from biotic sources. Biogenic mixing may be of particular importance in mangrove ecosystems, as these habitats experience comparatively little physical mixing due wind or tidal flow (8, 9).

Previous studies have explored the role of biogenic mixing contributions of several taxa, including pelagic jellyfish, fish, and krill in the water column (10, 11), as well as benthic bivalves (12). These studies suggest the potential for non-trivial levels of animal-mediated mixing even in unsheltered environments, although others suggest a much lower level of mixing contribution (13) in the global oceans relative to processes such as internal waves. Little work is available regarding such effects in shallow (< 2m) habitats, however. One process by which internal waves might be produced is through interactions between tidal flow and topography in stratified waters (14). However, in the shallow, quiescent conditions of mangrove lagoons, these conditions do not appear to be present.

While the primary source of turbulent mixing in shallow coastal ecosystems is wind-driven wave action (15), mangrove ecosystems provide shelter from wind and waves (9), as well as dampen mixing from high-velocity tidal currents, which tend to run primarily in deeper channels within mangrove ecosystems (16). Therefore, physical mixing is naturally low in *Cassiopea* sp. habitat, implying that animal-mediated mixing could have more substantial impacts in these environments.

The upside-down jellyfish, *Cassiopea* sp., has a nearly circumtropical distribution (17). They are found in shallow, low energy coastal environments, often dominated by the red mangrove, *Rhizophora mangle* (18) and have been previously documented at densities up to 31 animals m^−2^ or 20% benthic coverage (19). In our field site they were encountered at water depths of < 2 m. Their natural range is extensive, and is further spreading as an invasive species due to anthropogenic introduction (20–22) and through natural range expansion due to rising ocean temperatures (21). In addition, the size and abundance of *Cassiopea* sp. have increased near anthropogenically disturbed habitats, where nutrients tend to be elevated (23).

Unlike most other jellyfish, *Cassiopea* sp. exhibit an epibenthic lifestyle, with their bell on the substrate and oral arms facing upwards. They pulse their bells nearly continuously producing a vertical jet (24–26), however the extent of this vertical jet and volumetric flux have not been fully quantified. The direction of flow and volumetric fluxes may be particularly noteworthy given the fact that in the absence of *Cassiopea* sp. there is a downward flux of nutrients with the sediments acting as a nutrient sink in sheltered mangrove habitats (27). Thus, an upward pump at the benthos may serve to alter this flux and also interact with interstitial porewater, pulling a fraction of this water into the water column (26) and potentially altering local productivity.

In this study, we quantify the large-scale flow features of individual *Cassiopea* sp. as well as flows created by small groups both in the laboratory and in the field. We discuss the results in the context of environmental mixing in mangrove ecosystems and examine the role of biological pumping at the surface of the benthos with respect to the liberation of interstitial porewater from the sediments.

## Results

### Field Population

At our study site in the Florida Keys, USA, the density of the studied population of *Cassiopea* sp. ranged from 3 to 97 animals m^−2^, with a median density of 29 animals m^−2^ (fig. 2c). The sizes of these animals followed a Poisson distribution (fig. S1, Chi-Squared Test for Independence, p < .001) with a mean of 11.3 cm oral arm diameter (OAD) and 8.6 cm resting bell diameter (RBD), and there is a linear relationship between OAD and RBD (fig. S2, eq. 1, n = 50, R^2^ = 0.9). We calculated that animals with the observed average RBD of 8.6 cm have a wet weight of 56.5 g (fig. S2, eq. 2, n = 50, R^2^ = 0.99) and a dry weight of 3.2 g (fig. S2, eq. 3, n = 31, R^2^ = 0.98).

**Figure 1:**
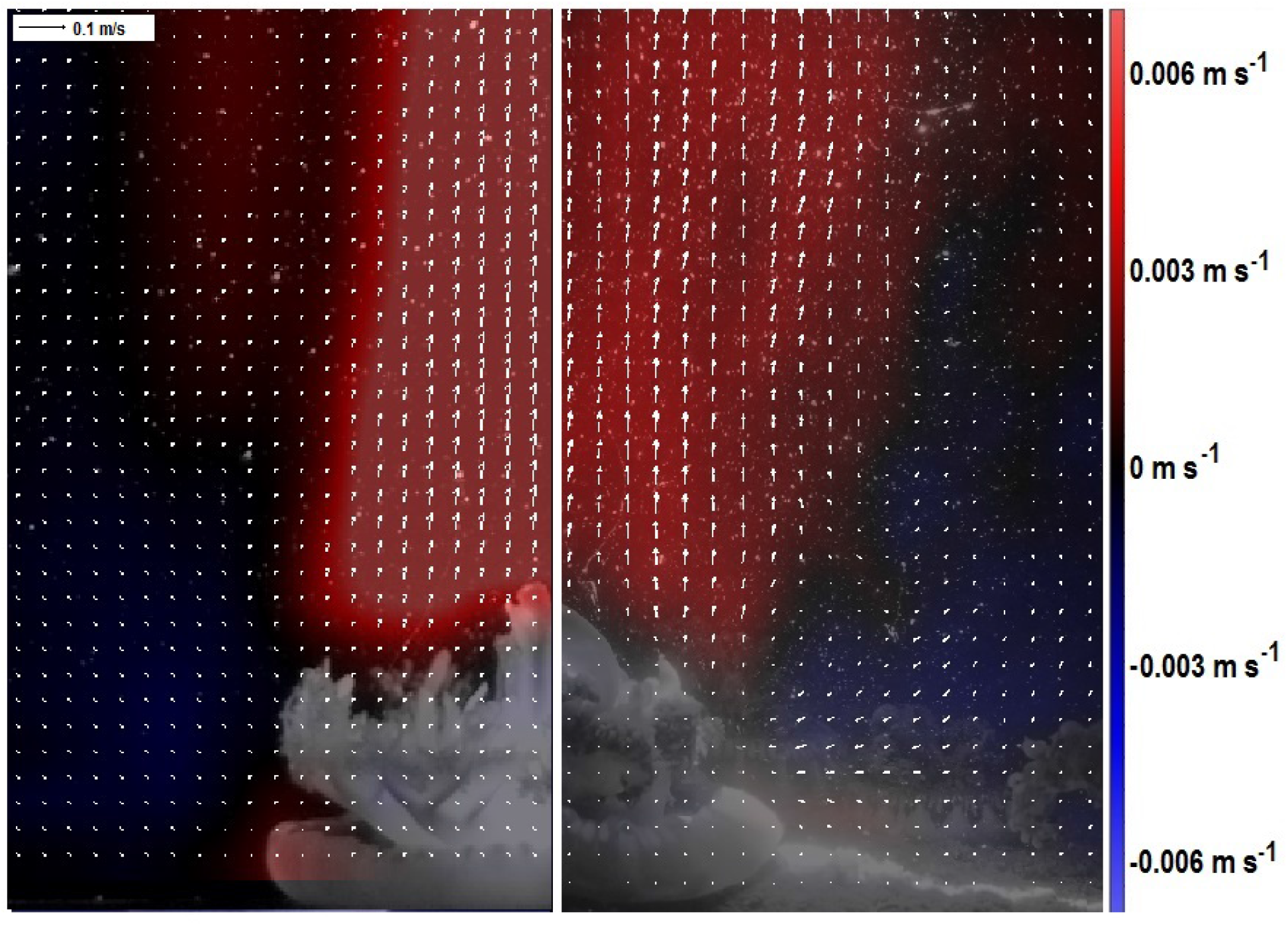
Laboratory (left) and in-situ (right) Particle Image Velocimetry (PIV) of a representative *Cassiopea* sp. jet. White arrows represent the velocity vectors while colored areas represent the flow with either an upward (red) or downward (blue) component.

**Figure 2:**
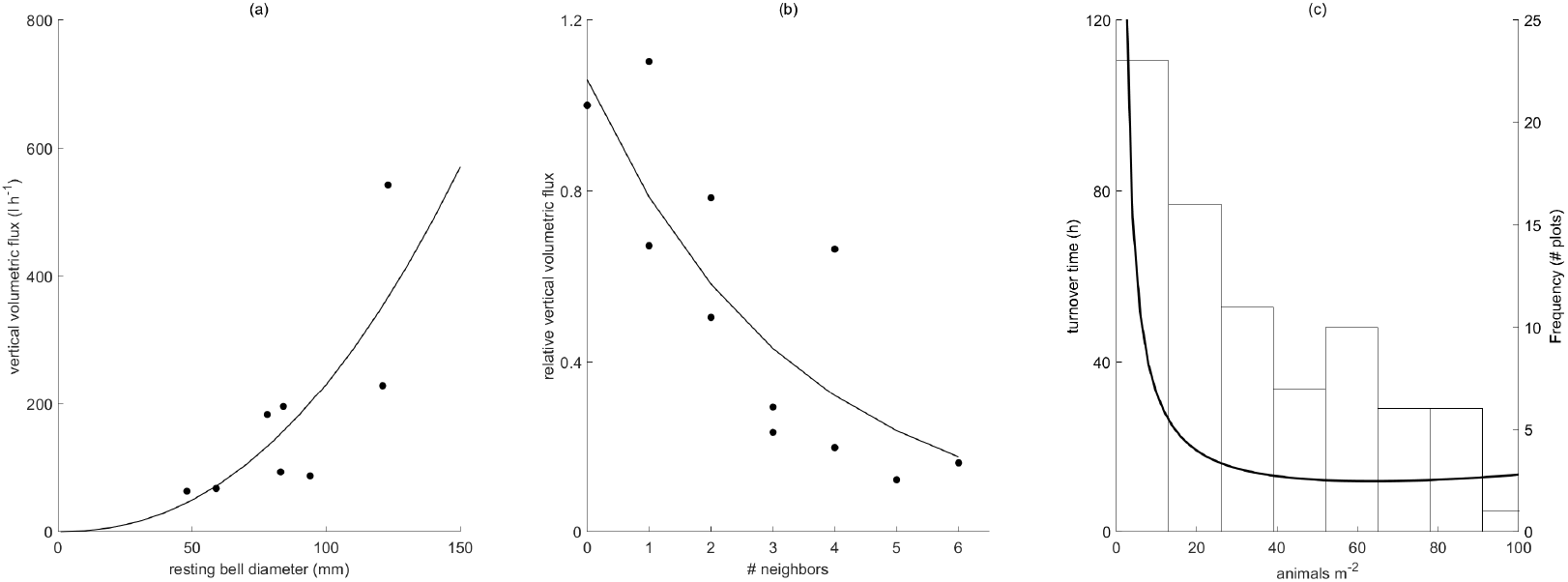
**(a)** Vertical flux of water increases exponentially and follows a power law relationship with animal size (R^2^ = 0.732, n = 9). **(b)** The relative vertical flux, such that a value of one is equal to the amount an individual pumps in the absence of neighbors, decreases as the number of neighbors increases (R^2^ = 0.74, n = 12). **(c)** Predicted time to turn over the column of water above a square-meter patch of *Cassiopea* sp. assuming a water depth of 1 meter, which was typical in our field site, relative to the population density in that patch. Frequency of different population densities observed is shown as a histogram of 1 m square plots.

### Quantitative Description of the Vertical Jet

Particle Image Velocimetry (PIV) was performed on 9 *Cassiopea* sp. ranging in size from 4.8 cm to 14.5 cm resting bell diameter (RBD). Analysis of the shape of the jet produced by these animals showed that the jet diameter (D_j_), relative to RBD, spreads linearly as height (Z) increases, (fig. S2, eq. 4, Linear regression: *R*^2^ = 0.64, *p* < 0.001). Maximum velocity at each height correlated by a power function to height normalized to bell diameter (fig. S2, eq. 5, *R*^2^ = 0.33). Our regression predicts a mean velocity of 20 mm s^−1^ at benthos level regardless of animal size, but the rate of water speed slowing above this height decreased as animal size increased. At a height of one bell diameter above the animal, *Re* = 1091 based on jet diameter. An extrapolation assuming calm conditions in the field shows that for the average-sized animal (RBD = 8.6 cm) the jet would slow to 3.86 mm s^−1^ at a height of 2 meters. Since *Cassiopea* sp. in this study were found only between 0.5 and 2 m deep, it is assumed that the jets will reach the surface in their natural habitat. In-situ PIV confirmed that wild *Cassiopea* sp. in the field produced maximum vertical flow velocities similar to our laboratory-based velocity measurements.

### Flux and Turnover Time

At a relative distance of two bell diameters above the animals, vertical velocity was found to be independent of bell diameter (Spearman’s rank correlation, *ρ* = 0.233, *p* = 0.551), as was expected from the observation of jet shape. Because jet diameter was found to correlate directly with bell diameter (Spearman’s rank correlation, *ρ* = 0.9, *p* = 0.002), and velocity was independent of bell diameter, vertical flux of water (Q) was found to follow a power law function relative to resting bell diameter, with a power of 2.23 (fig. 2a; fig. S2, eq.6, n = 9, R^2^ = 0.74). The relationship can also be expressed as weight-specific flux (F) in terms of dry weight (fig. S2, eq. 7). Thus, an average-sized *Cassiopea* sp. moves 212 l h^−1^ of seawater, a mass-specific volumetric flow rate of 62.9 l h^−1^ g^−1^ dry weight (DW).

Since *Cassiopea* sp. form aggregations in nature, a correction factor was needed to account for flow-related interactions between individuals. The jet of an individual *Cassiopea* sp. was examined under conditions with different numbers of neighbors, and the flux was found to decrease with the addition of each neighbor (fig. 2b; fig. S2, eq. 8), where neighbors were defined as being in physical contact with each other. Based on photographic transects of the field population in the Florida Keys, at the median density of 29 animals m^−2^, each animal will have an average of 1.23 neighbors (fig. S2, eq. 9). Assuming a water depth of 1 meter and a vertical jet that reaches the surface, the turnover time for the water column was calculated. A single animal would turn over the water column over a square meter of benthos in 4.7 hours. At the observed median population of 29 animals m^−2^ the resulting flux is 3980 l h^−1^, and the calculated turnover time is reduced to 15 minutes. Turnover time decreases as population density increases, to a minimum of 11.8 minutes at a density of 64 animals m^−2^ (Fig. 2c) at which point turnover time begins to increase due to increased interference between neighbors.

### In Situ Flow Measurement

In the Lido Key site, in-situ PIV demonstrated that *Cassiopea* sp. increase vertical transport of water in these habitats. Mean vertical velocity increased from 0.12 mm s^−1^ downward (± 0.39 mm s^−1^ SD, n = 7) in the absence of jellyfish to 4.01 mm s^−1^ upward (± 1.52 mm s^−1^ SD, n = 5) (Fig. 3). This difference was determined by an unpaired T-test to be highly significant (T = 5.8487, df = 10, p = 0.0002). The horizontal component of the flow showed a smaller, but still significant change, reducing from 19.82 mm s^−1^in the absence of jellyfish to 12.97 mm s^−1^when a jellyfish was present (T = 3.2021, df = 10, p = 0.0095). Turbulent kinetic energy dissipation rates were 3.5 × 10^−6^ m^2^s^−3^ (± 1.8 × 10^−6^ m^2^s^−3^ SD, n = 4). In the presence of jellyfish this increased to 6.5 × 10^−6^ m^2^s^−3^ (± 4.3 × 10^−6^ m^2^s^−3^ SD, n = 8). This increase, while measureable, was not statistically significant (Two-sample T-Test, p = 0.2).

**Figure 3:**
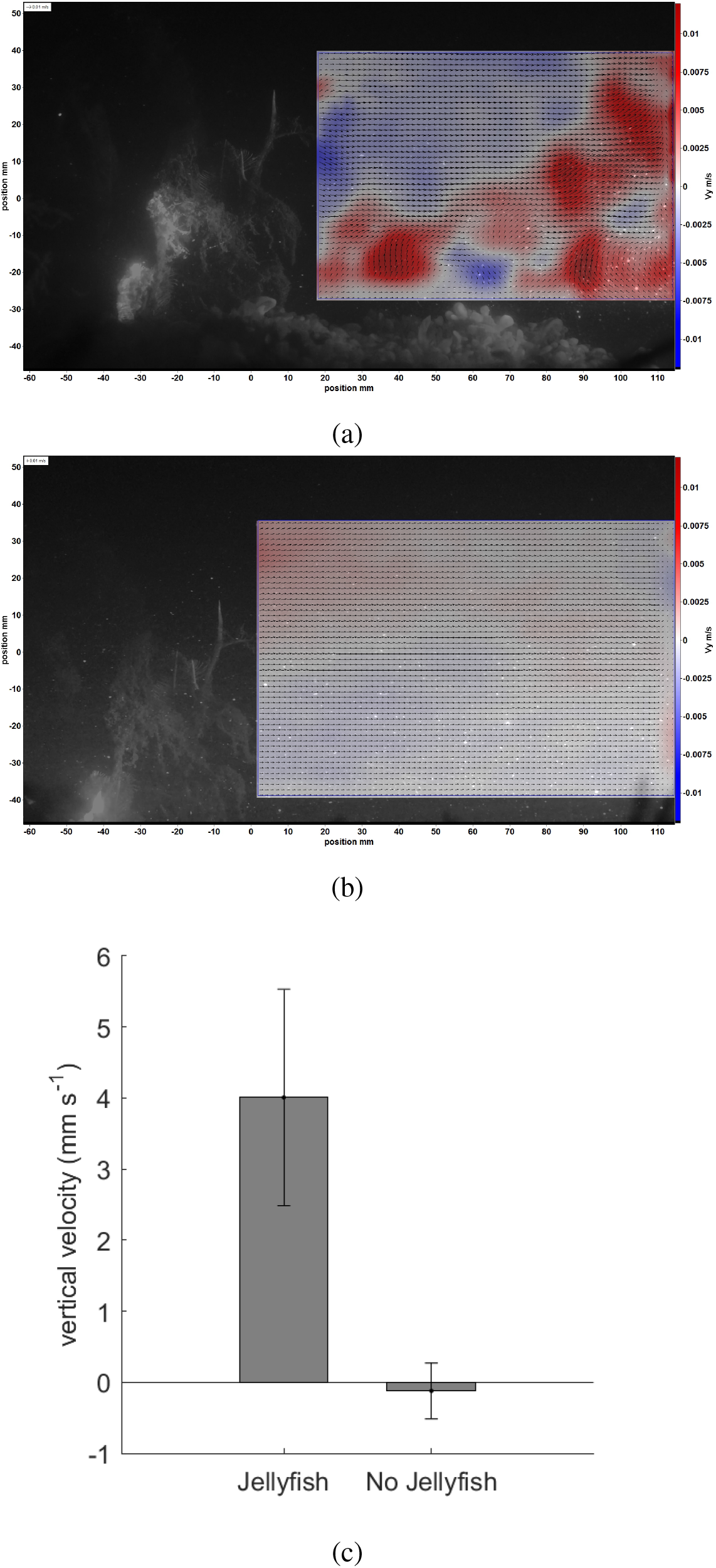
Representative time-averaged vertical flow component in the presence (a) and absence (b) of *Cassiopea* sp. at Lido Key, Florida as measured using particle image velocimetry over 1 second. Blue indicates downward flow, red indicates upward flow. Masked area was unavailable to PIV analysis due to the presence of solid structure, including the jellyfish and surrounding habitat structure. Quantitative analysis (c) shows that the presence of *Cassiopea* sp. significantly increased vertical flow from −0.12 cm s^−1^ to 4.01 cm s^−1^. (Unpaired t-test, df = 10, *p* = 0.0002)

To confirm that these habitats are quiescent and protected from wind-driven wave action, a comparison between wind waves between the mangrove habitat and the adjacent Sarasota Bay showed a reduction in wave height from 5.4 ± 2.4 cm (N = 10) in the bay to 0.07 ± 0.07 cm (N = 10) in the mangroves (Supplemental Materials). In addition, the horizontal velocity measured during our PIV experiment was 19.82 cm s^−1^ in the absence of jellyfish. Since this experiment was performed near peak tidal flow in order to avoid resampling water masses, this value must be near the upper limit for tidal velocities in this habitat.

### Interstitial Water Release

Interstitial water release was quantified by measuring changes in concentrations of fluorescein in water over fluorescein-laced sand. Fluorescein dye concentrations increased visibly in the trials that included a jellyfish but not in controls. The ratio of absorbance at 494 nm between t=3 h and t=2 h increased from 1.02 (± 0.07 SD, n = 5) to 1.15 (± 0.16 SD, n = 9) in the presence of a jellyfish. From this absorbance we calculated the average flow attributed to a single jellyfish, normalized to an RBD of 8.6 cm, which amounted to 2.64 ml h^−1^ (± 2.1 ml h^−1^ SD, n = 9) of interstitial water released into the water column (Fig. 4). Trials without jellyfish had no apparent fluorescein release. The flow rate in the presence of jellyfish was found to vary significantly from zero (1-Sample t-test, p = 0.0055). Interstitial water release rate increased linearly with bell diameter. (Fig. 4, n = 9, R^2^ = 0.3622).

**Figure 4:**
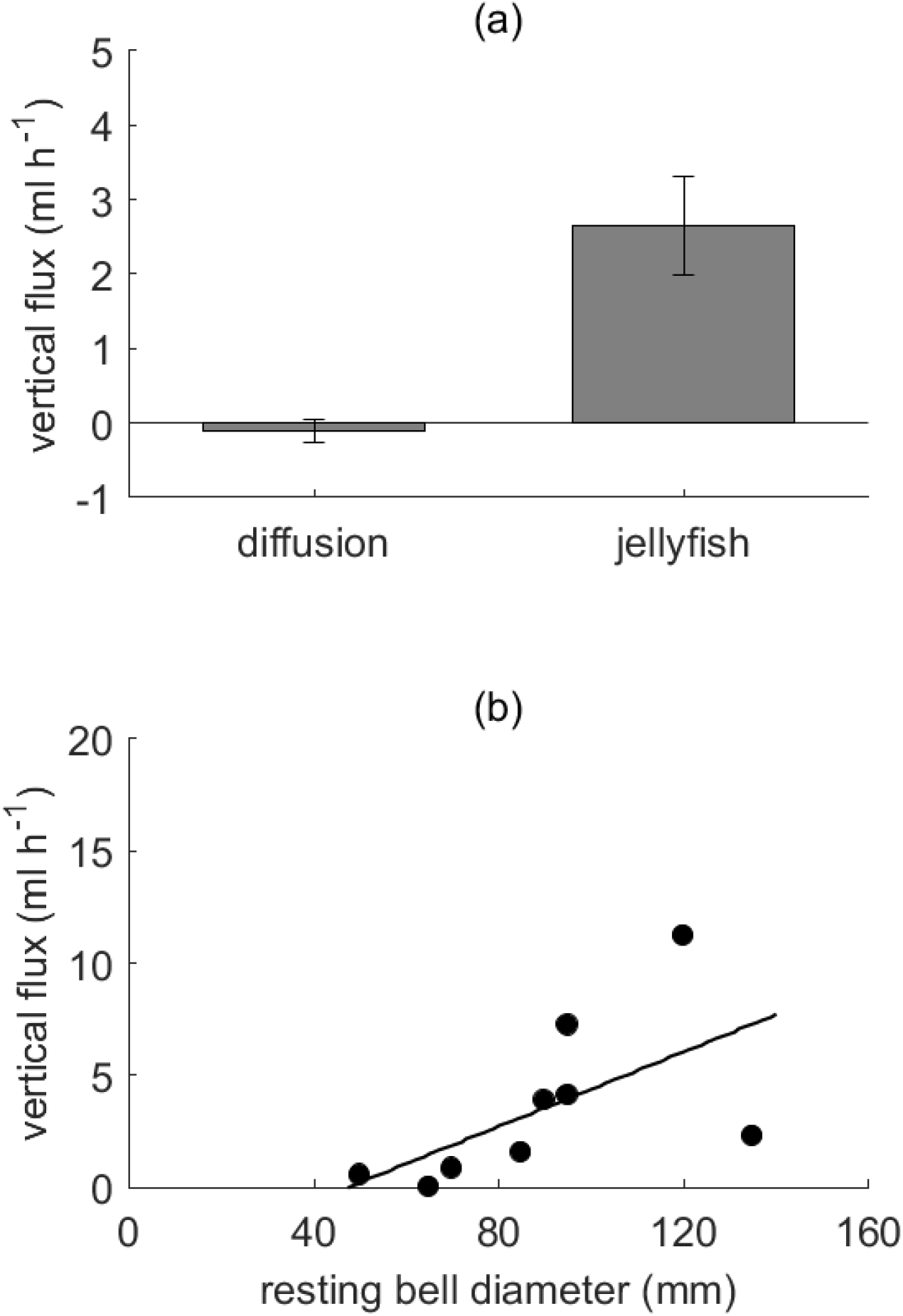
**(a)** Vertical flux of interstitial water out of the sediment attributed to a single *Cassiopea* sp., normalized to a resting bell diameter of 8.6 cm (the average observed in the field), compared to the rate due to diffusion in the experimental container. Interstitial water release due to diffusion was negligible, while the average *Cassiopea* sp. released 2.6 ml h^−1^. **(b)** Vertical flux of interstitial water relative to bell diameter. As animal size increased, the interstitial flow increased as well (n = 9, R^2^ = 0.3622).

## Discussion

In many coastal ecosystems, wind and tidal flow are the primary mixing forces (15). However, the ecosystems inhabited by *Cassiopea* sp. are sheltered by dense stands of *Rhizophora* mangrove trees, reducing wind-water interaction in these lagoons and estuaries (9). In addition, *Cassiopea* sp. dwell outside of the tidal channels, where most tidal flushing in these habitats occurs (8), residing instead on the adjacent flats, at depths of < 2 m. Together, these two habitat features result in a system with comparatively little physical mixing. Further evidence of the quiescent nature of this habitat is given by the presence of a fine-grain benthic substrate in the areas where *Cassiopea* sp. are abundant (35). Our results from field measurements near *Cassiopea* sp. habitats found turbulent kinetic energy dissipation rates comparable to previously reported measurements (36) from mangrove habitats confirming that environmental flows in which *Cassiopea* sp. are found are low, with little turbulent mixing. As such, the cumulative effect of a population of *Cassiopea* sp. has the potential to provide a large contribution to mixing, relative to non-biogenic factors, which would have important ecological implications.

Our measurements and calculations show that the average animal has the potential to produce a jet on the order of several meters tall, which would certainly affect the entire depth of the water column (0.5 m to 2 m) in the shallow ecosystem where *Cassiopea* sp. occur. This flow occurs at transitional Reynold’s numbers, and this is reflected in the slowing rate of the vertical jet. Our results indicate that an individual average-sized animal can move 212 l h^−1^ of seawater upward in its vertical jet (Fig. 2a) at velocities on a similar order of magnitude to those found in prior work (25). After factoring in the effect of neighboring animals, our model predicts that the median population density observed in our field site (29 animals m^−2^) can turn over a 1 m deep column every 15 minutes (fig. 2c). This biogenic contribution to environmental mixing is higher than any other known epibenthic species (Table 1).

**Table 1:**
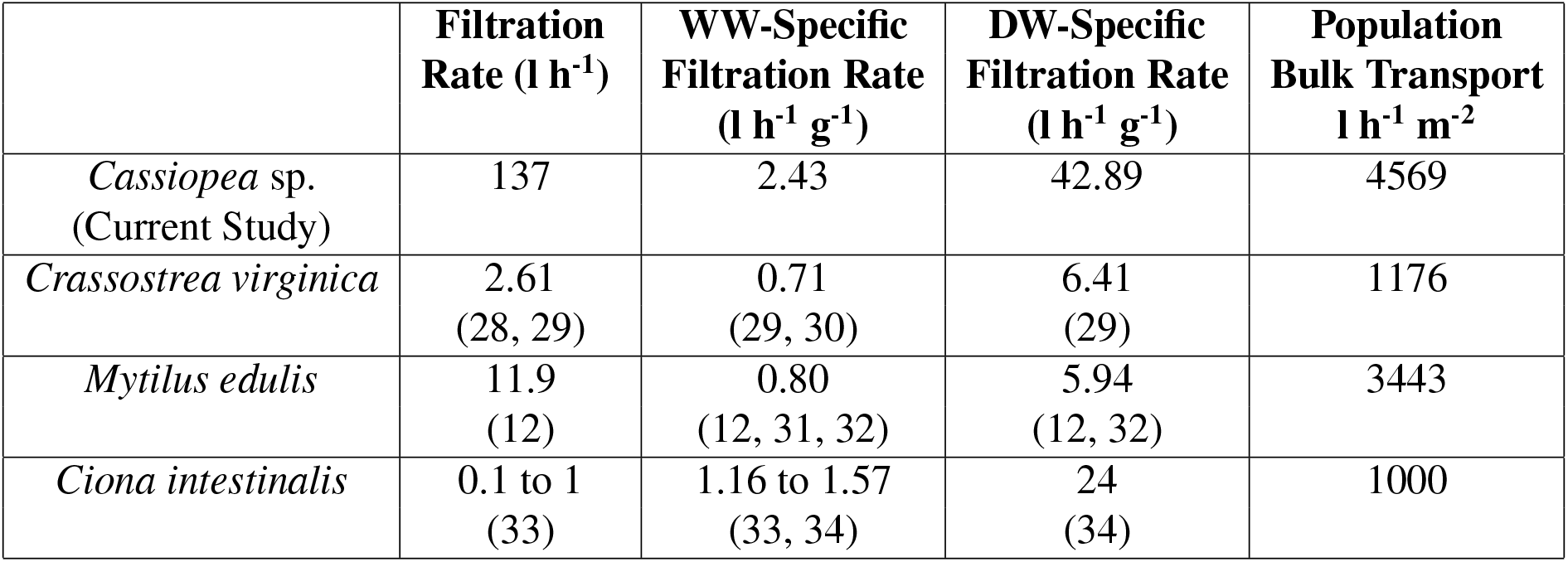
Estimated filtration rate per individual, as well as wet weight and dry-weight specific filtration rates for *Cassiopea* sp. (assuming a density of 29 animals m^−2^), *Crassostrea virginica*, *Mytilus edulis*, and *Cionia intestinalis*. Note: Rates for *Cassiopea* sp. are consistently an order of magnitude larger than those of bivalves. Citations refer to sources of filtration rates and size conversion factors used to calculate these values.

For comparison (Table 1), the mussel *Mytilus edulis* has been observed to pump 6.4 - 11.9 l h^−1^ and the oyster *Crassostrea virginica* has been calculated to pump 2.61 l h^−1^, in both cases two orders of magnitude below the rates measured in *Cassiopea* sp. in this study. To account for differences in body mass and body water content, dry weight-specific filtration rates can also be compared (Table 1). The eastern oyster *Crassostrea virginica* moves 6.4 l h^−1^ g^−1^ dry weight and *M. edulis* pumps 5.94 6.4 l h^−1^ g^−1^ dry weight. By comparison, *Cassiopea* sp. exhibit volumetric flow rates an order of magnitude higher than these bivalves by dry weight. Therefore, per unit of carbon, *Cassiopea* sp. act as much more effective water pumps than do bivalves. In terms of wet weight, *Cassiopea* sp. have weight-specific filtration rates that are also an order of magnitude above those of bivalves (Table 1). The weight specific pumping rate of tunicates is at a level closer to that of *Cassiopea* sp.. However, due to the much smaller size of *C. intestinalis*, they move a lower magnitude of water than *Cassiopea* sp. overall.

Of course, animals such as oysters and mussels can occur at much higher densities than *Cassiopea* sp. do, but their relatively small individual flow is not fully compensated for in terms of numbers. High densities of over 700 animals m^−2^ have been reported for both *Mytilus edulis* (37) and *Crassostrea virginica* (28). Assuming adult (0.3g dry weight (28)) oysters and additive contributions (a likely overestimate) this would produce approximately 1176 l h^−1^ m^−2^ flow. For mussels (32) and the same assumptions, we calculate a flow of 3443 l h^−1^ m^−2^. The tunicate *Ciona intestinalis* was found to pump water at rates of 0.1 to 1 l h^−1^ (33). This species, has been recorded at densities of over 1000 individuals m^−2^ (38) have a population filtration rate on the order of 1000 l h^−1^ m^−2^. Our median population of *Cassiopea* sp. (29 animals m^−2^) moved 3980 l h^−1^ m^−2^, while peak densities observed in this study moved 4569 l h^−1^ m^−2^. This translates to peak bulk flow estimates from *Cassiopea* sp. that are 36% higher than peak values for mussels, 386% higher than oysters, and 457% higher than benthic tunicates. Thus, *Cassiopea* sp. populations, despite their lower population densities, are responsible for greater amounts of water transport than are bivalve reefs or tunicate colonies and is occurring in an area with comparatively little physical mixing, in comparison to *M. edulis* which inhabits much more physically dynamic conditions (39). In addition, *Cassiopea* sp., unlike other biological pumps that are smaller, are capable of producing jets which reach the surface of the water (fig. S2, eq. 5). Other, smaller pumps are much more limited in the extent of their influence, with *M. edulis* jets decomposing within 5 cm of the bottom (40). The result of this difference in extent is that *Cassiopea* sp. is able to turn over the entire water column, while bivalves and other smaller organisms mix only the lower strata.

The fact that median *Cassiopea* sp. populations can turn over the entire shallow water column in 15 minutes has implications for several ecosystem processes. An increase in mixing, especially in upward vertical transport, could lead to a decrease in sedimentation in low-energy habitats. Particulates in mangrove habitats flocculate, increasing particle size, but the low density of these flocs reduces sedimentation rate (41). *Cassiopea* sp. themselves may contribute to flocculation by the production of mucus (19), which traps suspended particles (41). On the other hand, these particles may be further prevented from settling by addition of mixing energy from *Cassiopea* sp. thereby increasing the export of nutrients from mangrove ecosystems to neighboring systems such as coral reefs and seagrass beds. Further investigations would be required to determine the effects of *Cassiopea* sp. on sediment transport. Another important consideration for *Cassiopea* sp. mediated biogenic mixing is that the strongest flows are generated near the benthos. This is in contrast to wind-driven mixing in which the strongest flows are at the surface and may have important implications for productively in these habitats. For example, a mesocosm study where mixing was initiated close to the bottom, as is the case with *Cassiopea* sp., found that faster turnover resulted in longer-lasting algae blooms and higher chlorophyll concentrations than at lower mixing rates (6), likely because of the increased availability of nutrients from the sediment to phytoplankton cells. In addition, the accelerated exchange of water near the benthos to the surface in the presence of *Cassiopea* sp. may affect oxygen and CO_2_ exchange rates at the surface, affecting carbonate pH buffering, respiration rates, and primary productivity throughout the water column.

It has been previously demonstrated that *Cassiopea* sp. can transport interstitial porewater in sediments from several cm deep in the benthos to the water column (26) and while the maximum velocity of this flow was similar to those in this study at 2 cm h^−1^, volume fluxes were not measured and thus the potential impact on the local ecosystem is unknown. In Gulf of Mexico sediments, increasing the advective benthic flow produced an increase in denitrification rates (42) so it is possible that in areas where *Cassiopea* sp. is abundant, it may also contribute to increased denitrification. Additionally, increased flow alters the redox gradient and microbial diversity in benthic sediments (43). Our results show that while over the course of the experiment (3 hours), diffusion released negligible amounts of benthic porewater, an average-sized animal (8.6 cm) would release approximately 2.6 ml h^−1^ of porewater (fig. 4), and the median population of 29 animals m^−2^ would therefore release 1800 ml day^−1^m^−2^. Given that these animals tend to move slowly over the bottom (Durieux et al. unpublished data), thereby encountering new sediments regularly, and the reported phosphate and nitrogen concentrations in the sediment (44) at Long Key, Florida, the action of *Cassiopea* sp. could release roughly 0.34 mg day^−1^ m^−2^ of NO_3_, 2.35 mg day^−1^ m^−2^ of NH_4_, and 1.03 mg day^−1^ m^−2^ of PO_4_ at median population densities. Since these released nutrients will be mixed into the water column upon becoming entrained in the vertical jets and assuming a 1 m water depth, this represents an increase of 29% in water column NH_4_, a 6% increase in PO_4_, and a 0.5% increase in NO_3_ relative to reported water column nutrient concentrations from these habitats (44). The addition of the mixing contribution produced by *Cassiopea* sp. into a habitat with low environmental water mixing, as well as a substantial release of interstitial water leading to an additional source of nutrients in this ecosystem, make it likely that *Cassiopea* sp. are previously unrecognized ecosystem engineers within mangrove habitats.

## Materials and Methods

### Field Population

The field study site on Long Key, Florida, USA (24.527 N,-80.814 W) was sheltered by red mangroves (*Rhizophora mangle*) and held a population of *Cassiopea* sp. at depths ranging from to 2 m deep. The benthic habitat where *Cassiopea* sp. were most abundant consisted of very fine sediment, surrounded by seagrass (*Thalassia* and *Syringopodium*).

The population structure of a natural *Cassiopea* sp. population was quantified during December 2016, by setting two transects through the patch, 10 meters apart, parallel to one another in a roughly east-west orientation. Quadrats of 1 m^2^ were sampled every 1.5 m on each transect to reduce investigator bias in selecting animals for measurement. Each quadrat was photographed from above, and the oral arm diameter (OAD) of each animal was measured from these images by measuring the longest observable distance from one oral arm tip to another oral arm tip. During the summer of 2017, fifty animals were collected and imaged from above and from the side. The resting bell diameter (RBD, used for lateral imaging) was measured during the resting phase between bell contractions using ImageJ image analysis software (45, 46), and was correlated to the OAD, wet weight (WW), and dry weight (DW) by linear regression to allow conversion between measurements. Based on these parameters, populations of *Cassiopea* sp. at different densities were calculated.

### Vertical Jet Imaging

To determine the volumetric flow rate of vertical water movement, the vertical jet must first be described quantitatively. *Cassiopea* sp. of various sizes were collected from the Keys Marine Laboratory on Long Key, Florida, USA during the month of December, 2016 and August, 2017, and transported in collected seawater back to the University of South Florida, in Tampa, Florida, where they were housed in a 300 liter closed loop aquarium system. The animals were kept in artificial seawater mixed to a salinity of 35 with Instant Ocean aquarium salt, over a substrate of aragonite sand and high intensity metal halide lighting on a 12:12 light cycle.

The imaging setup consisted of a 45 × 45 × 45 cm aquarium, filled with artificial seawater. The water was seeded using 10 *μ*m reflective hollow glass spheres for particle image velocimetry (PIV). An Edgertronic high-speed camera filming at 50 frames per second provided a field of view ca. 30 cm x 30 cm. Two 2-watt continuous wave DPSS lasers (wavelength = 520 nm), spread through cylindrical lenses to produce narrow light sheets, were staggered one above the other to illuminate a single coronal plane across the entire field of view. One jellyfish at a time (n = 9) was placed on the bottom in the center of the aquarium, such that the laser sheet crossed the center of the animal. After allowing it to settle for about 10 minutes, 30 seconds of video were recorded at 50 frames per second. At least five such image sequences were made, and the three with the most similar pulse rates were retained for analysis using the LaVision software package, producing a PIV time-average over the 30 seconds. PIV was processed with interrogation windows between 48 and 64 pixels, at 50% overlap.

A custom MATLAB script was used to identify and measure the vertical jet from this time average. The location of the jet was defined as the region where the vertical velocity was greater than 0.5 mm s^−1^. This region was used to calculate the jet diameter. Due to the asymmetric meandering of the jet, volumetric flux was calculated by using a variant of the washer method. We integrated vertical velocity between the edges of the jet, using half-washers on each side of the position of maximum upward velocity. Because of the high degree of irregularity in *Cassiopea* sp. jets, the three image sequences of each animal were combined to create a single representative jet by taking the median values for jet diameter, jet area, maximum and average Vz, and vertical volumetric flux (Q) at each height for use in further analysis.

This method produces an artificial tapering of the jet shape towards its highest end due to the fixed velocity threshold, since the maximum velocity of the jet approaches the minimum threshold as it slows with increasing height. To account for this, data above the height of the maximum measured jet diameter were excluded. In addition, the development region of a turbulent jet does not follow the same patterns as the fully developed region (47). To take this into account, the lowest data points were excluded so that the data series always began at a local maximum in Vz.

For confirmation of the laboratory results, in-situ PIV was performed at the field site. A Nikon D750 DSLR at 50 frames per second camera in an underwater housing was lined up the bottom near an individual *Cassiopea* sp. with a green laser, spread through a cylindrical lens, also housed in a water-tight case. PIV analysis using the naturally-occurring particles was performed using DaVis, and the time-averaged vertical velocities were compared to those measured in the laboratory experiment.

### Quantitative Description of the Vertical Jet

Upward velocity (Vz) was expected to show an inverse relationship (47) with height (Z) relative to resting bell diameter (RBD) (fig. S2, eq. 10). In addition, the jet diameter (D_j_) relative to RBD was hypothesized to increase linearly over this region (47) (fig. S2, eg. 11). Vertical volumetric flux (Q) was expected to increase linearly with height, due to entrainment of water into the turbulent jet, and increase linearly with animal size (47). The resulting equation is a function of average velocity and jet diameter(fig. S2, eq. 12). These relationship were tested using the Nonlinear Regression tool in SPSS statistical software version 23.

To confirm that *Cassiopea* sp. in the field produce the same vertical jet as animals in a captive setting, in-situ PIV was performed on an animal at the Keys Marine lab, and the parameters of this jet were compared to those captured in the laboratory.

Weight-specific volumetric flow rates were calculated by calculating the expected flux of the animals used for morphological measurements, and then expressing this flux in terms of the wet and dry weights of those animals.

### Interactions Between Multiple Animals

Because *Cassiopea* sp. are often found in dense aggregations, we determined the degree of interference between adjacent animals using the same PIV design. A single animal was placed in the filming vessel, and then surrounded by an increasing number of neighbors, each in direct contact with the bell of the study animal. The vertical velocity and diameter of the original animal’s jet were calculated as before, allowing us to determine the degree of inhibition caused by the addition of neighbors. This was compared to the number of neighbors expected in wild populations, determined from the same transect photos used to determine average animal size and population density.

### Turnover Time

The time needed for a *Cassiopea* sp. population to turn over the water column above it, where the jet reached the surface, was calculated by taking the volume of water above a 1 m^2^ patch of a hypothetical *Cassiopea* sp. population and calculating the vertical volumetric flux for each animal in that population. The sum of the vertical flux rates (Q) of the animals in this hypothetical patch is divided by the volume of water, giving the turnover rate (fig. S2, eq. 13).

### In Situ Flow Measurement

Flow velocities were measured around a population of *Cassiopea* sp. at Lido Key, Florida, (27.303 N, −82.566 W) using an in-situ PIV apparatus recording at 60 fps. The PIV plane was aligned with the direction of tidal flow, and the water column above and downstream of a single Cassiopea was recorded over a period of several minutes. The animal was then removed, and recording continued. After PIV vectors were computed, 10 second time-averages were taken of the vectors across the entire field of view, spaced apart temporally by a time period determined to allow the tidal current to carry the previously measured water mass out of the field of view. This was done to prevent re-sampling the same water mass. Turbulent Kinetic Energy (TKE) dissipation rates ca. 15 cm above the bottom near patches of *Cassiopea* sp. was were calculated from the PIV data using published methods (48), assuming homogeneous and isotropic conditions. Additional confirmatory methods using acoustic doppler velocimetry can be found in the supplemental materials.

### Interstitial Water Release Rates

We labeled play sand by mixing it with a solution of fluorescein in artificial seawater until the sand was uniformly damp. A smooth layer of labeled sand 2.5-3 cm deep was added to the bottom of an 8 l plastic bucket in a larger water bath, capped with a ca. 1 cm deep layer of clean play sand mixed with seawater at a rate of 500 ml seawater to 2 l of dry sand, which delayed leaching of fluorescein into the water column by about 30 minutes. The bucket was then filled with 4 liters of artificial seawater without disturbing the sand layers. Immediately after filling we took samples of interstitial and column water. A *Cassiopea* sp. specimen was then placed on the sand in the center of the bucket. On each day of experimentation, an additional trial was performed without a jellyfish to measure diffusion rates in the absence of animals. At two and three hours, the water column was sampled and the absorbance of this water at 494 nm was compared to a dilution curve of the interstitial water sample. The relative fluorescein concentrations at each timepoint were corrected by subtracting both the measured concentrations of the starting water sample for the same trial (to control for unintentional fluorescein released during setup), and the concentration of the control bucket at the same time point (to control for diffusion). The change in these corrected values between two and three hours was then used to calculate the rate of interstitial water release into the water column.

## Acknowledgments

The authors would like to acknowledge the following people who assisted with specimen collection and experimentation (in alphabetical order): Olivia Blondheim, Alicia Durieux, Christian Fender, Olivia Hawkins, Kathrene Lo, Gabrielle Scrogham, Nils Tack. Additionally, we would like to thank the Keys Marine Laboratory and Dr. David Lewis from the University of South Florida, for use of their facilities. Funding for this project was provided by the National Science Foundation (CBET-1511996, IDBR-1455471 and OCE-1829945 to B.J.G).

