## Supplemental Materials for "Benthic jellyfish dominate water mixing in mangrove ecosystems"

December 9, 2020

### Morphological Measurements of Cassiopea spp.

The standard methods of measuring jellyfish, such as wet weight and bell diameter, require manual handling of each animal. This limits their use in field surveys, and is inconvenient for behavioral experiments where physical manipulation might create methodological artifacts. For example, in our transect survey of animal sizes, photographs of quadrats were taken from above, and the oral arms often obscured the bell margins of the animals. Knowing the relationship between bell diameter and oral arm length was an essential component of this analysis.

#### Methods

We collected 50 *Cassiopea sp.* specimens of different sizes by hand from the Keys Marine Laboratory in Layton, Florida in the summer of 2017. These animals were placed in a 10-gallon aquarium and video-recorded next to a scale from above and from the side. The animal was then removed from the water, and the maximum diameter to which the bell could be spread was measured (Maximum Bell Diameter, MBD) to the nearest millimeter with a ruler. Each animal was gently shaken in air to remove water, and it was then weighed on a tared tray (Wet Weight, WW), following which the animal was frozen and returned to the University of South Florida. There, 31 of the specimens were dried for at least 48 hours in a drying oven and reweighed (Dry Weight, DW).

From the video, the bell diameter was measured from the side view during the resting phase between bell contractions (Resting Bell Diameter, RBD), and the maximum distance across the oral arms was measured from above (Oral Arm Diameter, OAD) using ImageJ image analysis software. The correlations between these measurements were calculated by linear regression.

There was a strong linear correlation between Maximum Bell Diameter and both Resting Bell Diameter (Eq. 1, $R^2 = 0.97$) and Oral Arm Diameter (Eq. 2, $R^2 = 0.94$). The relationship between Maximum Bell Diameter and both Wet Weight (Eq. 3, $R^2 = 1.00$) and Dry Weight (Eq. 4, $R^2 = 0.98$) followed a power function with an exponent near 3, as would be expected in a length-to-volume conversion.

$$\text{RBD}_{(mm)} = 0.846 * \text{MBD}_{(mm)} \tag{1}$$

$$\text{OAD}_{(mm)} = 1.23 * \text{MBD}_{(mm)} - 12.228 \tag{2}$$

$$\text{WW}_{(g)} = 3 * 10^{-5} * \text{MBD}_{(mm)}^{3.1039} \tag{3}$$

$$\text{DW}_{(g)} = 4 * 10^{-7} * \text{MBD}_{(mm)}^{3.3594} \tag{4}$$

### Population Structure in the Florida Keys

During December of 2016, two parallel transects were set 10 m apart across a patch of *Cassiopea sp.* at the Keys Marine Laboratory in Layton, Florida. To avoid sampling bias, a 1 m square quadrat was placed every 1.5 m along the transect, and was photographed from above. Using the quadrat as a scale, the OAD of each animal in the photographs was recorded, using ImageJ image analysis software. In addition, we counted the number of neighbors that each animal had, defined as overlap of the oral arms.

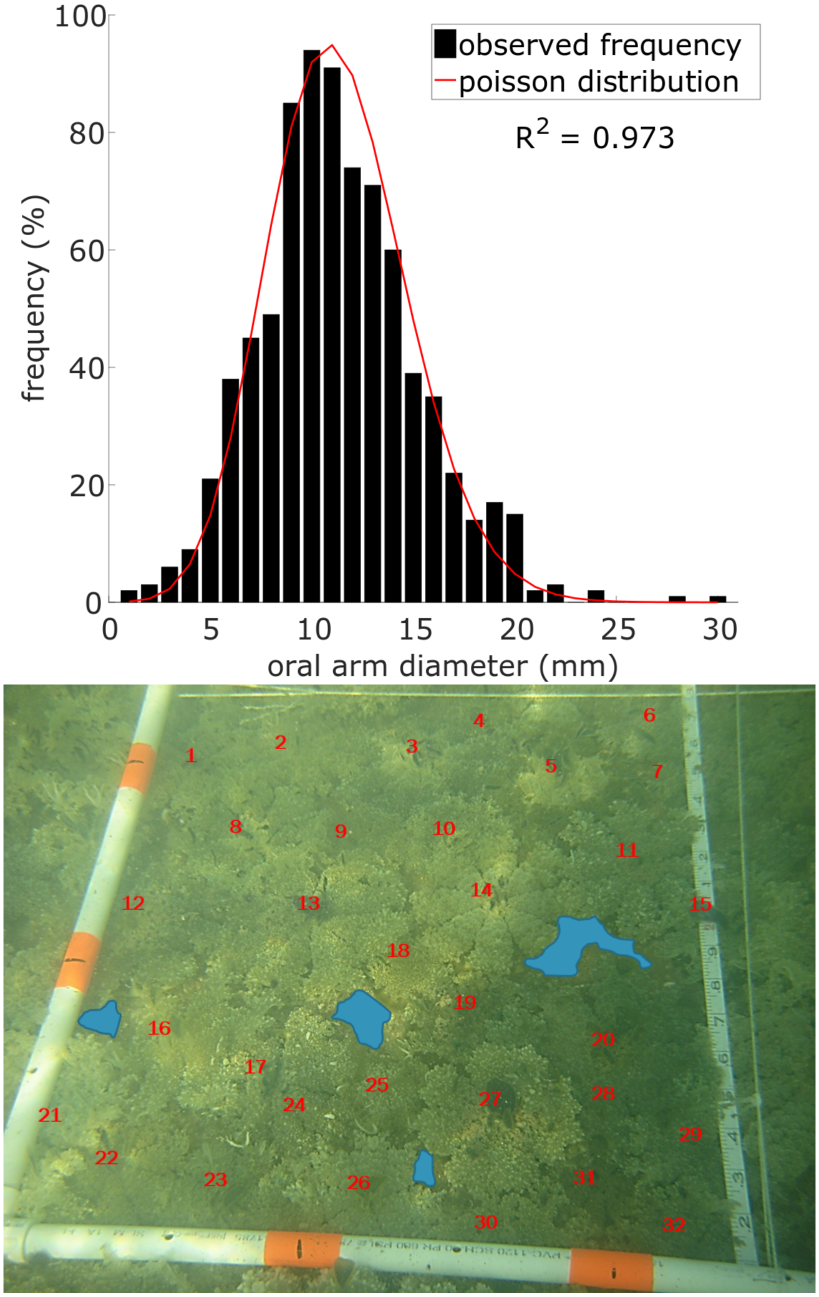


Figure 1: Top: Relative frequency of size classes of *Cassiopea sp.* (n = 799) from Long Key, Florida, follows a Poisson distribution (Chi-Squared test for independence, $p < .001$).
Bottom: A representative 50cm quadrat in a patch of high *Cassiopea sp.* population density at the Keys Marine Laboratory in Layton, Florida. Red numbers indicate individual *Cassiopea sp.*, and areas of exposed bottom were marked in blue.

#### In Situ Flow Measurement

In the Lido Key site, acoustic doppler velocimetry demonstrated that *Cassiopea sp.* increase vertical transport of water in these habitats. Mean vertical velocity increased from 2.1 mm $s^{-1}$ (±

Equations:

$$RBD_{\text{mm}} = 0.73 * OAD_{\text{mm}} \tag{1}$$

$$WW_{\text{g}} = 1 \times 10^{-4} * RBD_{\text{mm}}{}^{2.97} \tag{2}$$

$$DW_{\text{g}} = 2 \times 10^{-6} * RBD_{\text{mm}}{}^{3.21} \tag{3}$$

$$\frac{\text{D}_{\text{j}}}{RBD} = 0.49 * \left(\frac{Z}{RBD_{\text{mm}}}\right) + 0.59 \tag{4}$$

$$V_{\text{Z Max}} = 14.41 * \left(\frac{Z}{RBD_{\text{mm}}}\right)^{-0.41829} \tag{5}$$

$$Q_{\text{l h}^{-1}} = 0.008 * RBD_{\text{mm}}{}^{2.23} \tag{6}$$

$$F_{\text{l h}^{-1}\text{g}^{-1}} = 96.89 * DW_{\text{g}}{}^{0.68} \tag{7}$$

$$Q_{\text{rel}} = 1.5367e^{-0.35 * (\text{N} + 1)} \tag{8}$$

$$\text{Neighbors} = 0.033 * \text{Density}_{\text{m}^{-2}} + 0.267 \tag{9}$$

$$V_{\text{z}} = a * \left(\frac{Z}{RBD}\right)^{b} \tag{10}$$

$$\frac{\text{D}_{\text{j}}}{RBD} = a * \left(\frac{Z}{RBD}\right) + b \tag{11}$$

$$Q = V_{\text{Z(avg)}} * \pi * \left(\frac{D_{\text{j}}}{2}\right)^{2} \tag{12}$$

$$\text{Turnover Time}_{(h)} = \frac{\sum Q\left(\frac{m^3}{h}\right)}{\text{Depth}_{(m)} * 1\ m^2} \tag{13}$$

Figure 2: Equations referenced in the paper describing the morphology (1-3) biogenic flow (4-8), and population (9) by *Cassiopea sp.*, as well as the expected form of the fluid flow equations used in the paper (10-13).

1.4 mm $s^{-1}$ SD, n=7) in the absence of jellyfish to 3.3 mm $s^{-1}$ (± 2.25 mm $s^{-1}$ SD, n = 9) directly over patches of jellyfish (Fig. 3). This 57% increase in vertical flow represents a statistically significant difference (Unpaired T-test, $df = 14$, $p < 0.01$).

In order to verify that our study site on Lido Key, Florida, was sheltered from wind and wave effects, we used an anemometer to compare wind speed and wave height within the lagoon to the Western shore of Sarasota Bay, the windward side of Lido Key on the day of measurement. Wind speed was significantly lower in the lagoon ($0.65 \pm 0.64$ m $s^{-1}$) than the bay ($1.6 \pm 0.76$ m $s^{-1}$) (Two-sample T-Test, n = 10, p = 0.01). Wave height also varied significantly between the lagoon ($0.07 \pm 0.07$ cm) and the bay ($5.4 \pm 2.4$cm) (Two-Sample T-Test, n = 10, p < 0.0001). At that same time, the nearby COMPS-C21 Buoy (USF Coastal Ocean Monitoring and Prediction System) measured wind speeds averaging 7.8 m $s^{-1}$and a mean significant wave height of 25 cm.

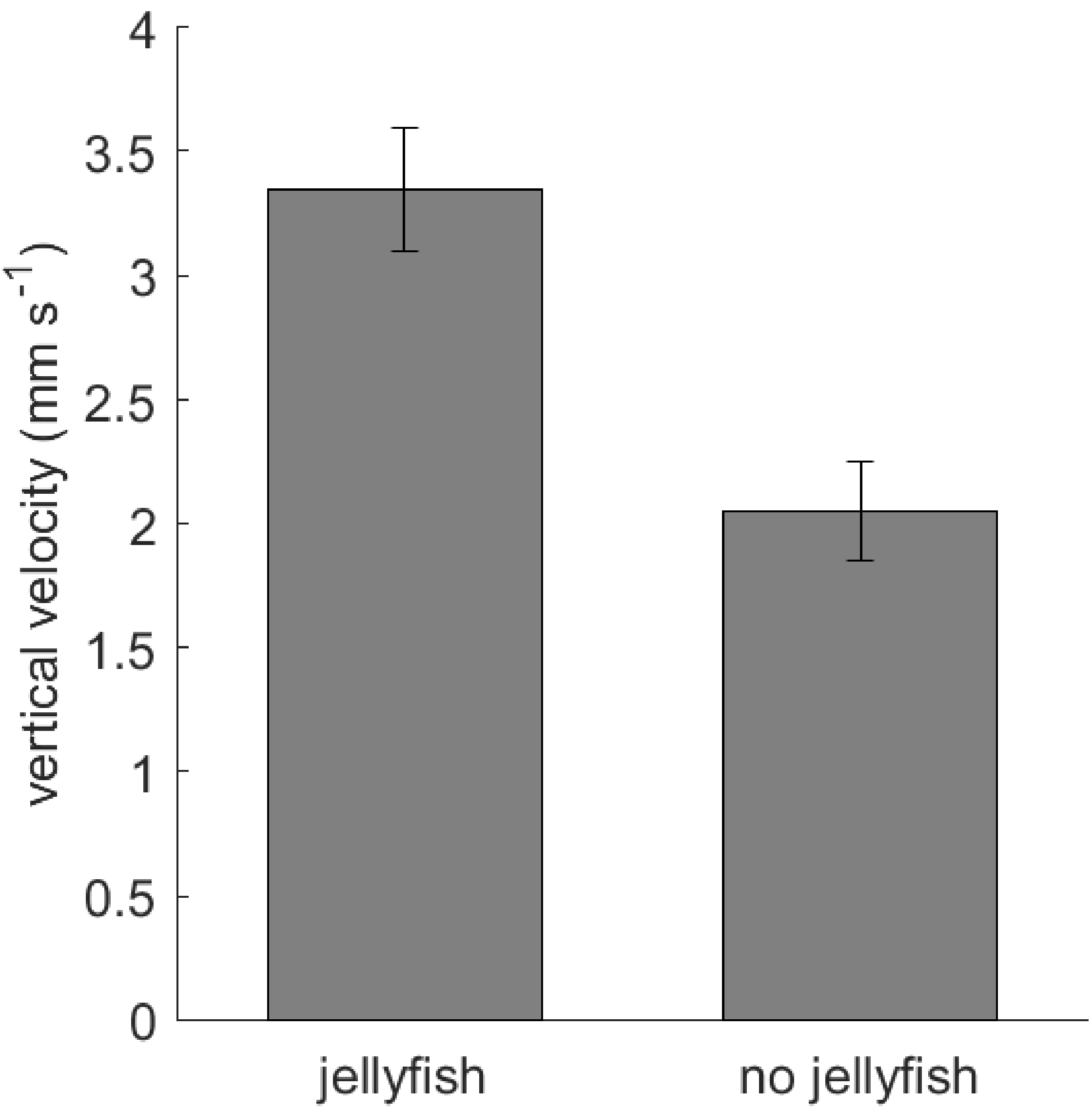


Figure 3: Vertical flow component in the presence and absence of *Cassiopea sp.* at Lido Key, Florida as measured using acoustic doppler velocimetry. The presence of *Cassiopea sp.* significantly increased vertical flow (Unpaired t-test, $p \leq 0.001$)
